# Local Topological Structures and Global Connectivity Patterns in a *Trachelospermum jasminoides*: a Pilot Study

**DOI:** 10.1101/2025.02.04.636494

**Authors:** Arturo Tozzi

## Abstract

Topological approaches to biological systems provide insights into their growth patterns, network connectivity and spatial organization. This perspective explores how biological structures self-organize, maintain stability and adapt to environmental constraints, revealing fundamental principles of efficiency, robustness, resilience and functional optimization. In this pilot study, we analysed the local and global topological properties of a *Trachelospermum jasminoides* bush (commonly known as star jasmine) using persistent homology, graph theory, spectral analysis and percolation theory. The spatial positions of individual flowers were extracted from an image of the bush and represented as a point cloud to capture their structural distribution and spatial relationships. Using Delaunay triangulation, a connectivity graph revealed a dominant connected component with minimal isolated structures. DBSCAN analysis identified a large number of small, localized clusters, reflecting biological and environmental influences. Most flowers connected to five to six neighbours, forming a uniform network with high clustering. Shortest path analysis showed efficient long-range connectivity, with paths avoiding sparse regions. Spectral analysis indicated smooth percolation without bottlenecks, while percolation analysis simulations revealed resilience up to 18% flower removal, after which connectivity broke down. In sum, we showed that the star jasmine bush topology balances local structural constraints with global connectivity, ensuring efficient resource distribution and structural integrity. By integrating topological data analysis with ecological modeling, we suggested a methodological approach to understanding natural growth networks. These insights can inform broader studies on biological pattern formation, network resilience and plant architecture modeling in ecology, agricultural sciences and biomimetic design.

## INTRODUCTION

The study of topological structures in biological systems provides a unique perspective on how organisms organize themselves in space, ensuring both functional efficiency and resilience. In plant systems, topology plays a critical role in determining how resources are distributed and how growth patterns emerge in response to environmental constraints (Li et al., 2017; Shimotohno and Scheres, 2019; Wang et al., 2020). The *Trachelospermum jasminoides* bush, with its intricate floral and vine structures, provides a good example for exploring these topological principles. By applying computational topology, graph theory and fractal growth models, we seeked to analyze both local and global spatial properties of a star jasmine bush, revealing the relationships arising from its structure. Traditional botanical research has long focused on morphological classifications, describing the physical characteristics of plants in terms of their leaves, stems and flowers. Still, advancements in mathematical and computational topology have opened new avenues for understanding plant growth beyond mere morphological description (Carlsson, 2009; Edelsbrunner and Harer, 2010; Boissonnat et al., 2018; Liu et al., 2020). The application of persistent homology, spectral graph analysis and percolation theory allowed researchers to uncover hidden structural patterns influencing the connectivity of biological networks (Robins, 1999; Cohen-Steiner et al., 2007; Bianconi and Ziff, 2018). By identifying long-range relationships, local clustering tendencies and the thresholds at which structural integrity is compromised, the self-organizing nature of plant growth can be better understood.

### Trachelospermum jasminoides

as a climbing plant, relies on flexible structural adaptations to balance between local constraints and global connectivity while maintaining an interconnected floral arrangement supporting pollination and resource transport. Our analysis explores the underlying principles shaping the topology of the star jasmine bush, focusing on how individual flowers distribute themselves and identifying the threshold at which connectivity within the floral network begins to fragment. To analyze the topological properties of *Trachelospermum jasminoides* bush, our pilot study first extracted flower positions from an image and represented them as a spatial point cloud. By applying Delaunay triangulation, we constructed a network that captured the connectivity between neighboring flowers. This representation allowed for the application of network theory to analyze the formation of clusters, the presence of long-range connectivity and the overall resilience of the floral structure. One key measure in our analysis was the percolation threshold, which indicates how the structure responds to the progressive removal of flowers. If a significant portion of the network remains connected despite a high percentage of removals, the structure can be considered robust.

In sum, our study integrated topological data analysis, network science and biological modeling to uncover the hidden structures within a *Trachelospermum jasminoides* bush. These findings provide a broader understanding of how plants self-organize and maintain connectivity under natural constraints, offering new perspectives in biological organization, ecological modeling and bio-inspired design.

## MATERIALS AND METHODS

We examined images of a Trachelospermum jasminoides bush, taken in June during peak blooming. Trachelospermum jasminoides, commonly known as star jasmine, is an evergreen, twining vine prized for its star-shaped white flowers (Zhao et al., 2017). It blooms profusely in late spring to early summer, attracting pollinators like bees and butterflies (Cai et al., 2024). The analysis of the *Trachelospermum jasminoides* bush topology was conducted through a structured sequence of computational and mathematical techniques aimed at extracting persistent homology, spatial relationships, connectivity structures and growth patterns from an image of the plant (Zomorodian and Carlsson, 2005; Niyogi, 2008; Edelsbrunner et al., 2002; Chazal and Michel, 2021). The study involved several key stages, beginning with image preprocessing and flower position extraction, followed by the construction of a topological representation through Delaunay triangulation and network analysis. These representations were then subjected to percolation simulations, spectral graph analysis and clustering algorithms, all of which contributed to a deeper understanding of the plant’s topological organization.

The first stage of the analysis involved preprocessing the image to extract relevant spatial information. The original image was converted to grayscale to remove any unnecessary color information and to simplify further processing. A thresholding technique was then applied to enhance the contrast between the flowers and the background. Since star jasmine flowers are generally brighter than their surroundings, a high-threshold binary transformation was used, which assigned white pixels to flower regions and black pixels to the rest of the image. To refine the extracted features, noise removal techniques, such as morphological opening and closing, were employed to eliminate small artifacts that were not part of the actual floral structures. Once a clean binary image was obtained, contour detection algorithms were applied to identify and isolate individual flowers. The centroid of each detected contour was computed, generating a point cloud representation of the flower positions within the *Trachelospermum jasminoides* bush.

Once extracted the spatial distribution of flowers, the next step involved constructing a mathematical representation of their connectivity using Delaunay triangulation (Song et al., 2021). This technique partitions the set of points into non-overlapping triangles, ensuring that no point falls within the circumcircle of any triangle, thereby providing an optimal way to connect neighboring flowers. The resulting triangulated structure captured the local connectivity patterns of the flowers, forming a foundation for network-based topological analysis. Each flower was treated as a node in a graph, while the edges of the triangulated structure represented potential paths of biological or structural influence. Following the construction of the connectivity graph, an extensive network analysis was performed to quantify the structural properties of the star jasmine bush. The graph was analyzed to determine key metrics such as the number of connected components, degree distributions and clustering coefficients. The occurrence of large-connected components indicated that most flowers belonged to a single dominant structure, while smaller, disconnected components suggested isolated floral clusters. Clustering analysis was performed to identify distinct floral groupings within the star jasmine bush. The DBSCAN (Density-Based Spatial Clustering of Applications with Noise) algorithm was chosen for this purpose because it is well-suited for identifying clusters of varying densities while distinguishing outliers. DBSCAN operates by grouping points that are closely packed together while marking points that do not belong to any cluster as noise (Sander et al., 1998; Zhang 2019). The algorithm required two parameters: the minimum number of points needed to form a cluster and the neighborhood radius defining which points are considered close to one another. Clustering coefficient analysis (van Diessen et al., 2014) provided insight into the tendency of flowers to form tightly packed groups, which could be indicative of growth constraints or ecological interactions.

Additionally, the average shortest path length and network diameter were computed to assess how efficiently one part of the floral network could be reached from another. To assess the resilience of the floral network, a percolation simulation was performed by progressively removing flowers from the structure and monitoring the effect on connectivity (Grimmett, 1999; Bianconi and Ziff, 2018). In each iteration, a random subset of nodes was removed and the size of the largest remaining connected component was recorded. This process continued until the network became fragmented, revealing the percolation threshold at which large-scale connectivity collapsed and subsequently the level of resilience. The network was further examined through spectral graph analysis, which involved computing the eigenvalues of the graph’s Laplacian matrix (Zhang 2011). The spectral properties of the graph provided insight into its structural stability and the presence of bottlenecks or weakly connected regions, making it possible to infer how information or resources might propagate through the *Trachelospermum jasminoides* bush. The smallest nonzero eigenvalue, known as the algebraic connectivity, was particularly useful in determining how well the network resisted disconnection (Bao et al., 2023). A high algebraic connectivity points towards a robust structure with minimal vulnerability to localized disruptions.

## RESULTS

The analysis of the *Trachelospermum jasminoides* bush’s topological structure revealed a complex interplay between local clustering and global connectivity, suggesting an optimized natural arrangement balancing resilience and efficiency. The spatial positions of individual flowers were used to construct a connectivity graph based on Delaunay triangulation, which provided a geometric foundation for analyzing the relationships between neighboring flowers (**Figure 1A**). We demonstrated that most flowers belonged to a single dominant connected component, with only a few isolated structures scattered throughout (**Figure 1A**). The degree distribution of the graph indicated that most flowers were connected to an average of five to six neighboring flowers, forming a relatively uniform network with no significant outliers. Further network analysis revealed a high clustering coefficient, indicating that flowers tended to form tightly packed groups rather than being randomly dispersed. The clustering analysis using DBSCAN identified 59 distinct clusters of flowers, highlighting localized structural groupings that may correspond to biological or structural growth clusters. This means that, while the overall structure remained connected, localized groupings of flowers emerged within the bush. (**Figure 1B**). This behavior is consistent with known biological principles, where plants optimize spatial efficiency to maximize exposure to light and pollination opportunities while minimizing energy expenditure.

**Figure 1.**
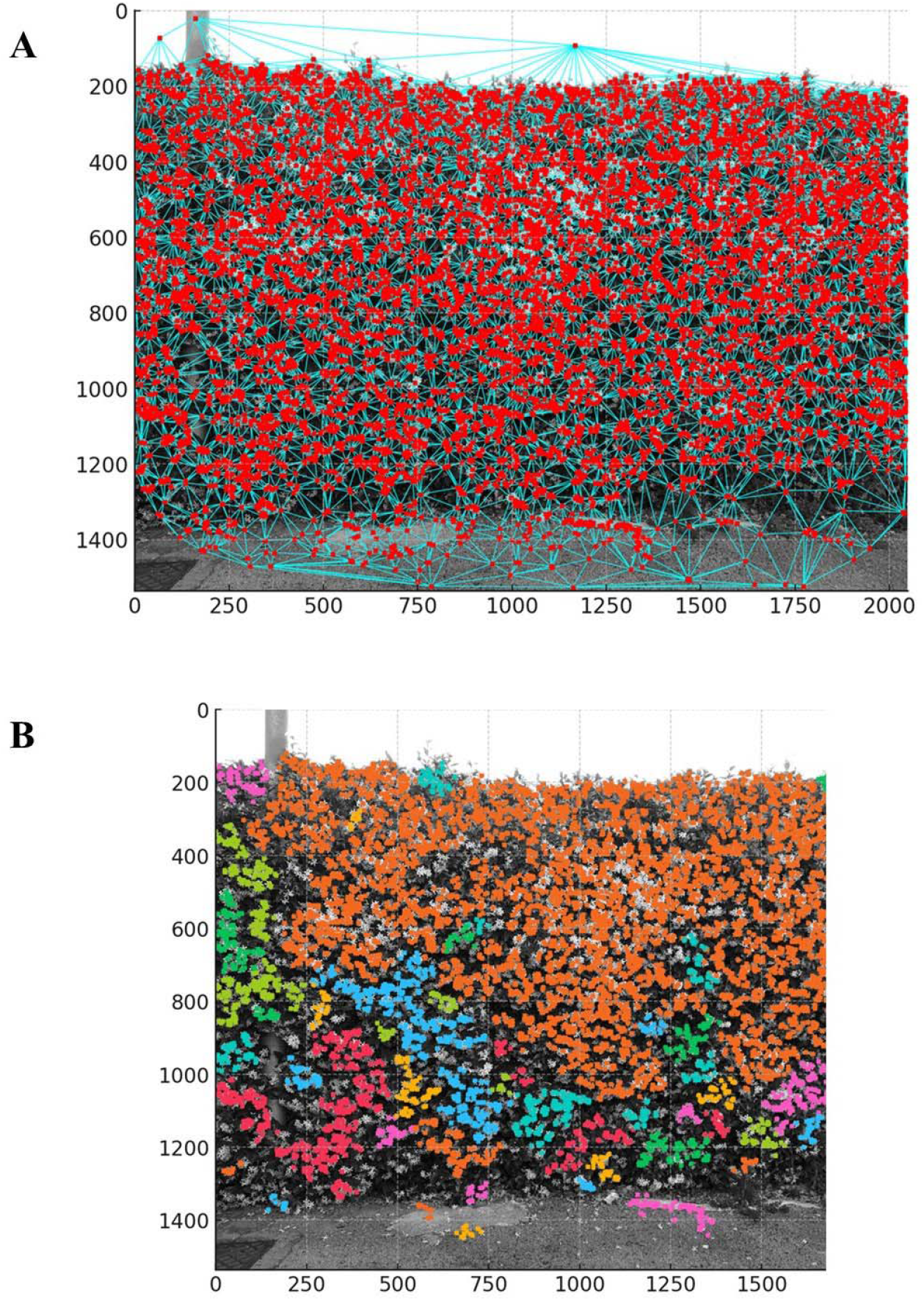
**A**. Delaunay triangulation highlighted short-range and mid-range connections between neighboring flowers. The flower centers were represented by red dots, marking individual flower positions. Some regions displayed higher connectivity, indicating clusters with strong local interactions. The largest connected component included 8,397 flowers, suggesting a globally connected, large-scale structure. **Figure 1B**. DBSCAN clustering analysis identified 59 distinct clusters of flowers. See text for further details.

The connectivity spread across the *Trachelospermum jasminoides* bush was further explored through the shortest path analysis. The minimal route between two distant flowers spanned 43 steps. This path followed the densest connectivity regions, demonstrating strong long-range percolation within the flower network. The start and end nodes were positioned in well-connected regions, effectively avoiding sparse areas, confirming the structural integrity of the network. Therefore, despite its dense clustering, the average path length between any two flowers remained relatively low. This suggests that the overall structure of the bush is optimized for both local interactions and long-range connectivity, ensuring efficient resource distribution and pollination. The percolation analysis provided additional insights into the robustness of the floral network. The star jasmine bush retained its large-scale connectivity until approximately eighteen percent of the flowers were removed (**Figure 2A**). This percolation threshold, relatively low compared to many random networks, highlighted the highly interconnected nature of the flower arrangement and indicated a high degree of resilience, as the structure remained intact despite significant disturbances. The rapid breakdown of connectivity beyond this threshold suggests that *Trachelospermum jasminoides* growth follows an optimized spatial strategy that prevents fragmentation under normal environmental conditions but becomes vulnerable when a critical mass of structural points is lost. Spectral analysis of the flower graph leveraged eigenvalues of the adjacency matrix in order to detect topological features such as bottlenecks, cycles and connectivity strength. This spectral analysis revealed a dense clustering of small eigenvalues, indicating strong overall connectivity within the star jasmine bush (**Figure 2B**). The smallest eigenvalues, ranging from 0.018 to 0.022, are characteristic of highly connected graphs with percolation-like behavior. The absence of significant bottlenecks suggested smooth percolation across long distances, reinforcing the idea that the network remained well-connected.

**Figure 2.**
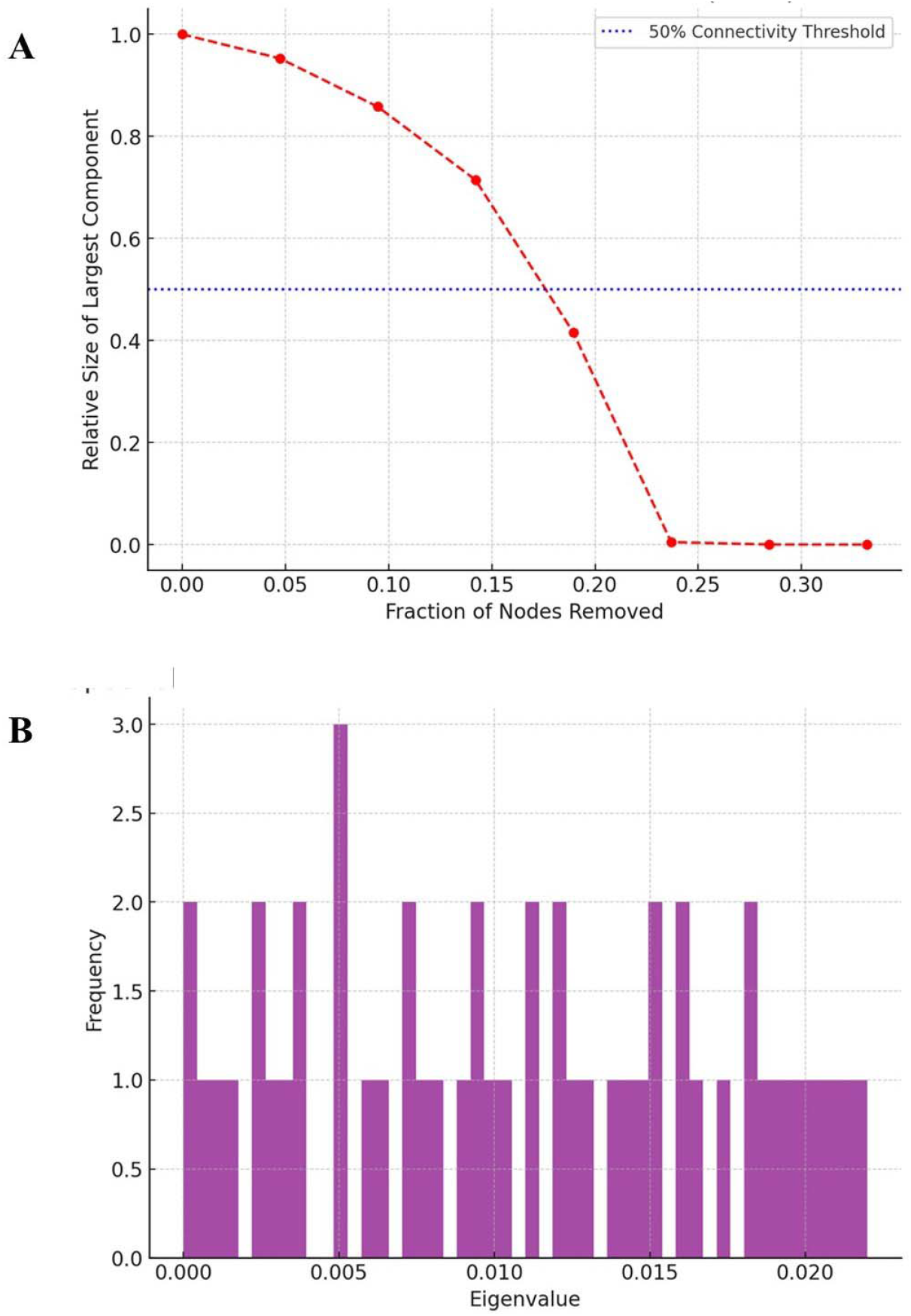
**A**. Percolation threshold simulation. Only 18.9% of flowers needed to be removed before the *Trachelospermum jasminoides* bush’s large-scale connectivity collapsed. The largest connected component remained stable up to approximately 15% removal, after which it rapidly disintegrated, indicating a sudden transition from a well-connected structure to fragmentation. **Figure 2B**. Spectral analysis leveraging the flower graph’s eigenvalue distribution of the adjacency matrix (first 50). A dense clustering of small eigenvalues was detected, indicating strong overall connectivity within the star jasmine bush. The smallest eigenvalues, ranging from 0.018 to 0.022, are typical of highly connected graphs with percolation-like behavior.

In sum, the presence of a smooth eigenvalue spectrum suggested that the connectivity of the floral network was evenly distributed without significant weak points or bottlenecks. Our findings support the hypothesis that the *Trachelospermum jasminoides* bush’s structural organization is not random but follows principles enhancing its overall integrity. The formation of localized clusters within a globally connected structure reflects an efficient organizational principle that maximizes functional interactions while maintaining large-scale stability.

## CONCLUSIONS

Using computational topology, graph theory and growth modeling, we provided an in-depth analysis of the topological properties of a *Trachelospermum jasminoides* bush. By extracting flower positions and analyzing their spatial relationships, we built a comprehensive model of the floral network, capturing both small-scale clustering behaviors and large-scale organizational patterns. Our results suggest that the star jasmine bush follow an optimized structural framework that maintain connectivity despite localized variations in floral density. One of the most significant findings was the discovery of a high clustering coefficient among the flowers, which suggested a strong tendency for localized grouping. This indicated that *Trachelospermum jasminoides* growth is not random but instead followed an underlying spatial organization to promote dense floral arrangements while maintaining connections to the larger plant structure. The clustering tendency aligned with known ecological and biological principles, particularly in relation to pollination strategies and resource distribution. Flowers arranged in tightly packed clusters may enhance pollination efficiency by increasing the likelihood that pollinators will move between closely spaced flowers before leaving the plant. This also has implications for the transport of nutrients and water, as connectivity within these clusters ensures effective internal resource distribution.

A key finding related to structural robustness emerged from the percolation analysis (Smirnov, 2001), which demonstrated that the *Trachelospermum jasminoides* bush maintains its connectivity even after the removal of approximately eighteen percent of its flowers. This suggests a high degree of resilience, as the plant is capable of withstanding significant structural changes without suffering immediate fragmentation. The observed percolation threshold aligns with results from similar analyses conducted in other biological networks, where plant and fungal structures exhibit similar levels of robustness. The rapid breakdown of connectivity beyond the eighteen percent threshold suggests that star jasmine plants have evolved an optimal distribution pattern that allows them to remain intact under typical environmental disturbances but may become vulnerable under extreme conditions such as disease outbreaks or mechanical damage. The spectral analysis of the star jasmine network reinforced these conclusions by providing insight into the plant’s global connectivity patterns. The eigenvalue distribution of the Laplacian matrix revealed a smooth spectrum indicative of a well-balanced and evenly connected structure. Supporting the findings from the percolation study, the presence of a high algebraic connectivity value confirmed that the *Trachelospermum jasminoides* bush is resilient to disconnection. The spectral properties of the network suggest that the star jasmine plant does not grow in an entirely uniform manner but instead follows a pattern of optimized connectivity, ensuring that floral structures remain accessible while avoiding excessive redundancy. The study’s spectral approach also allowed for the detection of subtle structural variations within the network that would have been difficult to observe using traditional morphological analysis alone.

When comparing these findings to other techniques commonly used in plant morphology studies, it becomes evident that computational topology provides unique advantages in uncovering hidden structural patterns. Traditional botanical studies rely on morphological classification systems that describe the arrangement of leaves, flowers and stems based on visual observation and measurement. While these methods are valuable for taxonomy and comparative morphology, they do not capture the deeper network properties influencing plant function and resilience. In turn, graph-based approaches offer a more quantitative framework for analyzing how plants organize themselves spatially. One of the primary advantages of our computational approach is its ability to model plant growth dynamically, providing testable hypotheses about the factors influencing *Trachelospermum jasminoides* topology. The presence of strong floral clustering raises questions about whether environmental variables such as sunlight availability, wind exposure or soil nutrient distribution contribute to the observed patterns. Future studies could test this hypothesis by conducting experiments in which star jasmine plants are grown under different environmental conditions to observe whether clustering behaviors change in response to external factors. Another testable hypothesis emerging from the percolation analysis is that star jasmine plants may prioritize connectivity to minimize the risk of fragmentation in response to mechanical stress. This could be experimentally verified by selectively pruning flowers from different parts of the plant and measuring how structural integrity is affected over time.

The findings of this study also have applications in ecological research, agriculture and biomimetic design. In ecology, understanding the topological organization of plants can provide insight into how different species compete for space and resources. The clustering behaviors observed in the *Trachelospermum jasminoides* bush could be compared to those of other climbing plants to determine whether similar patterns emerge in different species or whether star jasmine exhibits unique growth strategies. In agriculture, insights into plant topology can inform planting strategies to optimize space usage and resource efficiency. By applying principles of connectivity and clustering, farmers and horticulturalists could design planting arrangements that enhance pollination efficiency and nutrient transport while reducing vulnerability to disease spread. Biomimetic applications of plant topology extend into the realm of artificial network design. The ability of plants to maintain connectivity while minimizing redundancy offers valuable lessons for engineers developing self-organizing networks in telecommunications, transportation and material science. The star jasmine bush’s ability to remain robust under structural changes suggests that similar principles could be applied to the design of resilient infrastructure systems. Studying plant topology may also inspire new approaches to designing artificial materials that mimic the self-repairing and self-optimizing properties of biological systems.

Despite the strengths of this approach, certain limitations must be acknowledged. The study relied on a two-dimensional representation of the *Trachelospermum jasminoides* bush extracted from images, which may not fully capture the three-dimensional complexity of plant growth. While the extracted floral positions provided valuable insights into spatial organization, future studies could benefit from incorporating three-dimensional imaging techniques, such as LiDAR scanning or photogrammetry, to obtain a more complete structural representation. Additionally, the study focused primarily on flower distributions, without considering the role of underlying stem and leaf structures, which may also influence connectivity and resource transport. Including additional plant components in future analyses could provide a more holistic understanding of the plant topology. Incorporating biological signaling models alongside topological analysis could refine the accuracy of our findings. Additionally, the study did not explore temporal dynamics, meaning that it provided a snapshot of star jasmine topology rather than an analysis of how the structure evolves over time. Future research could address this by monitoring the growth of *Trachelospermum jasminoides* plants longitudinally, capturing real-time changes in connectivity and clustering.

The findings pave the way to further interdisciplinary research into the principles of plant topology, providing a framework for understanding how natural structures self-organize. The combination of computational topology, network science and biological modeling has demonstrated that plants operate under structural principles balancing local constraints with global connectivity. These principles appear to be fundamental across biological systems, influencing not only plant morphology but also ecological interactions, species competition and evolutionary strategies. By refining and expanding upon these methods, future studies can uncover the hidden structures governing plant growth, contributing to both theoretical biology and practical applications in ecology, agriculture and biomimicry.

## DECLARATIONS

### Ethics approval and consent to participate

This research does not contain any studies with human participants or animals performed by the Author.

### Consent for publication

The Author transfers all copyright ownership, in the event the work is published. The undersigned author warrants that the article is original, does not infringe on any copyright or other proprietary right of any third part, is not under consideration by another journal and has not been previously published.

### Availability of data and materials

all data and materials generated or analyzed during this study are included in the manuscript. The Author had full access to all the data in the study and take responsibility for the integrity of the data and the accuracy of the data analysis.

## Acknowledgements

none.

## Authors’ contributions

The Author performed: study concept and design, acquisition of data, analysis and interpretation of data, drafting of the manuscript, critical revision of the manuscript for important intellectual content, statistical analysis, obtained funding, administrative, technical and material support, study supervision.

## Declaration of generative AI and AI-assisted technologies in the writing process

During the preparation of this work, the author used ChatGPT to assist with data analysis and manuscript drafting. After using this tool, the author reviewed and edited the content as needed and takes full responsibility for the content of the publication.

